# Targeting AKT elicits tumor suppressive functions of FOXO transcription factors and GSK3 kinase in Multiple Myeloma

**DOI:** 10.1101/816694

**Authors:** Timon A. Bloedjes, Guus de Wilde, Chiel Maas, Eric E. Eldering, Richard J. Bende, Carel J.M. van Noesel, Steven T. Pals, Marcel Spaargaren, Jeroen E.J. Guikema

## Abstract

The phosphatidylinositide-3 kinases (PI3K) and the downstream mediator AKT drive survival and proliferation of multiple myeloma (MM) cells and several AKT inhibitors are currently being tested in clinical trials for MM patients. AKT inhibition has pleiotropic effects, and the key aspects that determine therapeutic efficacy are not fully clear. Therefore, we investigated the antimyeloma mechanism(s) of AKT inhibition. Among the various downstream AKT targets are Forkhead box O (FOXO) transcription factors, and we demonstrate that they are crucial for changes in gene expression upon AKT inhibition. Based on gene expression profiling we defined an AKT-induced FOXO-dependent gene set that has prognostic significance in a large cohort of MM patients, where low FOXO activity correlates with inferior survival. We show that cell cycle exit and cell death of MM cells after AKT inhibition required FOXO. In addition, glycogen synthase kinase 3 (GSK3), a negatively regulated AKT substrate, proved to be pivotal to induce cell death and to inhibit cell cycle progression after AKT inhibition. Finally, we demonstrate that FOXO and GSK3 induced cell death by increasing the turnover of the myeloid cell leukemia 1 (MCL1) protein. In concordance, the AKT inhibitor MK2206 greatly sensitized MM cells for the MCL1 inhibitor S63845. Thus, our results indicate that FOXO and GSK3 are crucial mediators of the antimyeloma effects of AKT inhibition, and suggest combination therapies that may have therapeutic potential in MM.

**KEYPOINTS:**

- FOXO transcription factors and the GSK3 kinase are pivotal tumor suppressors downstream of AKT inhibition in MM cells.
- FOXO and GSK3 activation after AKT inhibition leads to a decrease in MCL1 levels in MM cells resulting in cell death.

## INTRODUCTION

Multiple Myeloma (MM) is a malignancy of transformed clonal plasma cells that typically reside in the bone marrow. Despite considerable improvements in the median survival due to novel treatment modalities, patients inevitably relapse and become refractory to further treatment. Further understanding of MM and plasma cell biology is urgently needed and may lead to novel therapeutic strategies^1^.

The serine/threonine kinase AKT is a central node in the PI3K/AKT/mammalian target of rapamycin (mTOR) pathway, which is active in MM due to growth factors produced by the bone marrow microenvironment, or MM cells^2–5^. Furthermore, hemizygous deletions of phosphatase and tensin homolog (PTEN), a negative regulator of AKT, were reported in 5-20% of MM patients and human myeloma cell lines (HMCL)^6,7^. AKT signaling is involved in cell proliferation, survival and metabolism^3,8^. As such, it drives proliferation and sustains the increased energy requirement of MM cells by reprogramming various metabolic pathways^8^. Due to its crucial role in oncogenesis and cell survival, AKT is an attractive therapeutic target for various types of cancer including MM, and consequently, several clinical trials assessing the efficacy of AKT inhibitors in MM are ongoing^9^.

AKT has many substrates and pleiotropic effects in healthy and malignant cells. In addition to metabolic, translational and mitogen-activated protein kinase (MAPK) pathways^8^, forkhead box O transcription factors (FOXOs) and glycogen synthase kinase 3 (GSK3) are negatively regulated by AKT through phosphorylation^8^. The FOXOs, i.e. FOXO1, FOXO3, FOXO4 and FOXO6, are context-dependent transcription factors that act as tumor suppressors, but may also contribute to tumorigenesis^10^. Moreover, FOXO1 and FOXO3 have crucial and nonredundant functions in B-cell development, activation and differentiation^11–17^. FOXOs can be phosphorylated, acetylated and ubiquitinated by a wide range of enzymes, thereby regulating their stability, localization and activity^18^. Different interaction partners can also influence the specificity by which FOXO targets genes, regulating their expression^19^. AKT phosphorylates GSK3 on Ser9 (beta-isoform) and Ser21 (alpha-isoform), thereby inhibiting kinase activity^20–22^. GSK3 is a major AKT target involved in the regulation of cell death by controlling BCL2-family proteins^8,23–26^.

In light of the recent interest in AKT as a therapeutic target in MM, we set out to provide key insight into the antimyeloma mechanism(s) of AKT inhibition among its various downstream pathways. Here, we demonstrate that FOXO1/3 and GSK3 are AKT-restrained tumor suppressors, and that the expression of FOXO-dependent genes has prognostic value in a cohort of MM patients. Mechanistically, we provide evidence that the activation of FOXO and GSK3 provoked cell death in a nonredundant fashion through negative regulation of MCL1, a major anti-apoptotic protein in plasma cells and MM^26–28^. In accordance, AKT inhibition greatly sensitized MM cells for the MCL1 BH3-mimetic S63845, even in MM cells resistant to AKT inhibition alone.

Our results show that the antimyeloma effects of AKT inhibition hinges on the activation of FOXO1/3 and GSK3 and provide a clear rationale to explore combination therapies aimed at AKT and its downstream targets, such as MCL1.

## MATERIALS AND METHODS

### Cell culture and reagents

The human MM cell lines (HMCL) LME-1, MM1.S, XG-1, XG-3, LP-1, OPM-2, ANBL-6, UM-3, and RPMI-8226 were cultured in Iscove’s modified Dulbecco’s medium (IMDM; Invitrogen Life Technologies, Carlsbad, CA) supplemented with 2 mM of L-glutamine, 100 U/ml penicillin, 100 μg/ml streptomycin (Gibco, Thermo Fisher Scientific, Waltham, MA) and 10% fetal calf serum (FCS; Hyclone, GE Healthcare Life Sciences, Pittsburgh, PA). The cell lines XG-1, XG-3 and ANBL-6 were cultured in medium supplemented with 1 ng/ml interleukin-6 (IL-6; Prospec Inc, Rehovot, Israel), which was washed out prior to experiments. HEK293T cells were obtained from the American Type Culture Collection (ATCC, Manassas, VA) and cultured in supplemented Dulbecco’s modified eagle medium (DMEM; Invitrogen Life Technologies) and 10% FCS. The following small-molecule inhibitors were used: GSK2110813 (Afuresertib) (AKT inhibitor; Selleckchem, Houston, TX) MK2206 (AKT inhibitor; Selleckchem), CHIR99021 (GSK3 inhibitor, Sigma Aldrich, St. Louis, MO), AS1842856 (FOXO1 inhibitor, Merck, Darmstadt, Germany), S63845 (MCL1 inhibitor, Selleckchem), cycloheximide (Sigma Aldrich).

### Constructs and retroviral/lentiviral transductions

CRISPR/Cas9 knockout (KO) HMCL clones were generated by lentiviral transduction as described previously^29^. HMCLs overexpressing MCL1 were generated by retroviral transduction using the LZRS-*MCL1*-IRES-GFP plasmid. A more detailed description is available in the supplemental materials and methods.

### Patient samples

Primary tumor cells from MM patients (>80% plasma cells) were enriched by Ficoll-paque PLUS (GE Healthcare Life Sciences) density centrifugation. Patient material was obtained according to the ethical standards of our institutional medical ethical committee, as well as in agreement with the Helsinki Declaration of 1975, as revised in 1983. Primary MM cells were cultured overnight in IMDM + 10% FCS, supplemented with 1 ng/ml IL-6 before being used in further experiments.

### Immunoblotting

Immunoblotting experiments were performed as described previously^30^. Protocols and antibodies used are available in the supplemental materials and methods, densitometry quantification of immunoblots was performed using Image J software (imagej.net)^31^.

### Gene expression profiling

RNA from 2 × 10^6^ cells was isolated using TRI-reagent (Sigma-Aldrich) and purified using the RNEASY mini kit (Qiagen, Hilden, Germany) using the RNA cleanup protocol supplied by the manufacturer. The RNA was analyzed using Affymetrix Human Genome U133 Plus 2.0 arrays (Affymetrix, Santa Clara, CA) and normalized using MAS5.0 (accession no. GSE120941). Gene expression data was analyzed using the R2: Genomics Analysis and Visualization Platform (http://r2.amc.nl). Venn diagrams were prepared using BioVenn (www.biovenn.nl)^32^. Enrichment plots were generated using the Broad Institute gene set enrichment analysis (GSEA) computational method and software^33^.

### Flow cytometry

Specific cell death was assessed by 7-aminoactinomycin-D (7-AAD; Biolegend Inc, San Diego, CA) staining and flow-cytometry and calculated as reported earlier^34^. Cell cycle analysis was performed by determining DNA content and bromodeoxyuridine (BrdU) incorporation as described previously^29^. A detailed protocol is provided in the supplemental materials and methods.

### Statistics

Statistical analysis was performed using the Graphpad Prism software package (Graphpad Software, La Jolla, CA) and combination indexes were calculated using CompuSyn (ComboSyn Inc, Paramus, NJ)^35^. In case single dose drug combinations were used we calculated the expected effect of drug combinations (C) using the Bliss independence model [C=A+B-(A*B)] where A and B indicate the observed cell death at specific concentrations of the single drugs^36^.

## RESULTS

### Inhibition of AKT induces cell death in MM cell lines and patient samples

To investigate the effects of AKT inhibition we exposed human MM cell lines (HMCL) (n=9) to increasing concentrations of the allosteric AKT inhibitor MK2206 or the ATP competitive AKT inhibitor GSK2110183 (Afuresertib), both pan-AKT (AKT1/2/3) inhibitors that are currently being evaluated for clinical activity in MM and other cancer patients^37,38^. Both inhibitors potently induced cell death in the majority of the HMCLs. However, UM-3 and RPMI-8226 were refractory to AKT inhibitor-induced cell death **(Fig 1A)**. Enriched malignant plasma cells obtained from MM patients (n=6) **(Suppl Fig 1A)** showed a similar response, in which MK2206 induced cell death to a variable degree in all but one patient **(Fig 1B)**.

**FIGURE 1.**
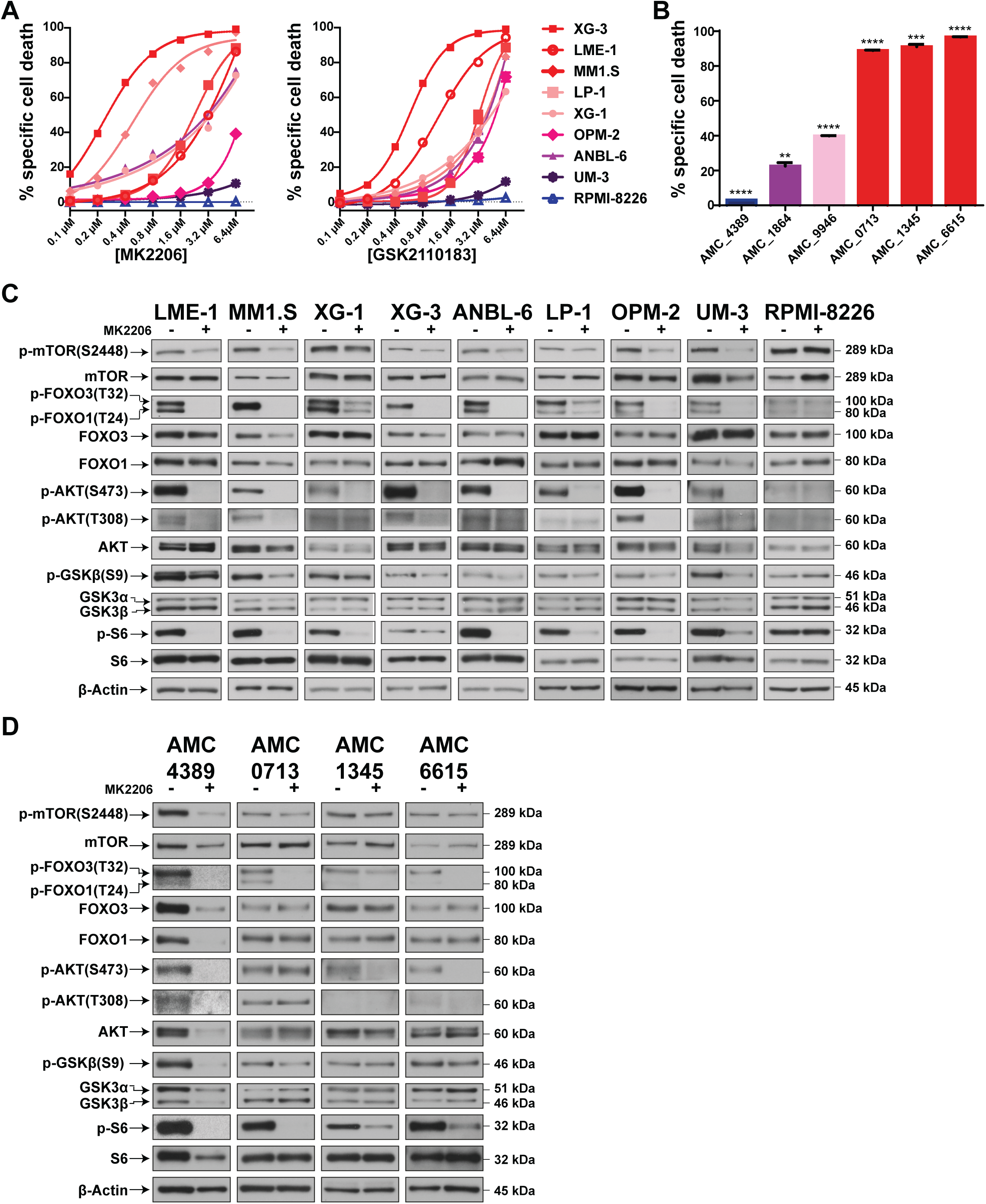
Inhibition of AKT in MM cells induces cell death. **(A)** Percent of specific cell death of HMCLs (n=9) treated with increasing concentrations of the ATP-competitive AKT inhibitor GSK2110183 (Afuresertib) (left panel) and the allosteric AKT inhibitor MK2206 (right panel) for 3 days. Specific cell death was calculated based on 7-AAD viability dye staining and flow-cytometry. Mean values of 3 independent experiments are shown. **(B)** Percent of specific cell death of primary MM plasma cells from patients (n=6) treated with 2.5 μM MK2206 AKT inhibitor for 3 days. Specific cell death was calculated based on 7-AAD viability dye staining and flow-cytometry. Means ± SEM of three technical replicates are displayed, n=3 (****p<0.0001; ***p<0.001; **p<0.01; one sample t-test). **(C)** Immunoblot analysis of protein expression in AKT-inhibitor treated HMCLs LME-1, MM1.S, XG-1, XG-3, ANBL-6, LP-1, OPM-2, UM-3 and RPMI-8226. Cells were serum starved for one hour, after which they were incubated in medium containing 10% FCS with or without 2.5 μM MK2206 for 2 hours. Shown are the indicated proteins, β-actin was used as a loading control. Representative immunoblot of at least 2 independent experiments is shown. **(D)** Immunoblot analysis of protein expression in primary MM patient plasma cells (n=4) serum starved for one hour, after which they were incubated in medium containing 10% FCS with or without 2.5 μM MK2206 for 2 hours. Shown are the indicated proteins, β-actin was used as a loading control.

To assess the downstream effects of AKT inhibition we performed immunoblotting for several established AKT targets. In all HMCLs except RPMI-8226, MK2206 decreased phosphorylation of AKT Thr308 and Ser473, targets of protein kinase PDK1 and mTOR complex 2 (mTORC2), respectively, that regulate AKT kinase activity^8^. The phosphorylation of downstream AKT substrates, FOXO1, FOXO3, GSK3beta and the ribosomal protein S6 were clearly decreased, indicating that MK2206 effectively blocked AKT function in all HMCLs except RPMI-8226. In agreement, similar effects of AKT inhibition were observed in primary MM patient samples **(Fig 1D)**.

These results confirm that FOXO and GSK3 are activated upon inhibition of AKT in the context of HMCLs and primary MM cells.

### FOXO transcription factors are required for AKT inhibitor-induced cell death of MM cells

To assess whether FOXO transcription factors were required for AKT inhibitor-induced cell death in MM cells we generated several FOXO1- and FOXO3-knockout clones for LME-1 (n=2), MM1.S (n=4) and XG-3 (n=4) using CRISPR/Cas9. Loss of FOXO protein expression was confirmed by immunoblotting **(Fig 2A)**. The loss of FOXO had no apparent effect on MM cell survival under basal condition (data not shown). However, AKT inhibitor-induced cell death was nearly abrogated in FOXO1-deficient LME-1 cells, whereas a small but significant reduction in cell death was observed in the FOXO3-deficient LME-1 cells **(Fig 2B, Suppl Fig 2A)**. In contrast, MK2206-induced and GSK211083-induced cell death was almost abolished in the FOXO3-deficient MM1.S and XG-3 cell lines, while FOXO1-deficient cells remained sensitive **(Fig 2B, Suppl Fig 2A)**. Of interest, we observed a slight increase in FOXO3 protein expression in FOXO1-deficient LME-1 cells, and a substantial increase in FOXO1 expression in FOXO3-deficient XG-3 cells **(Fig 2A)**.

**FIGURE 2.**
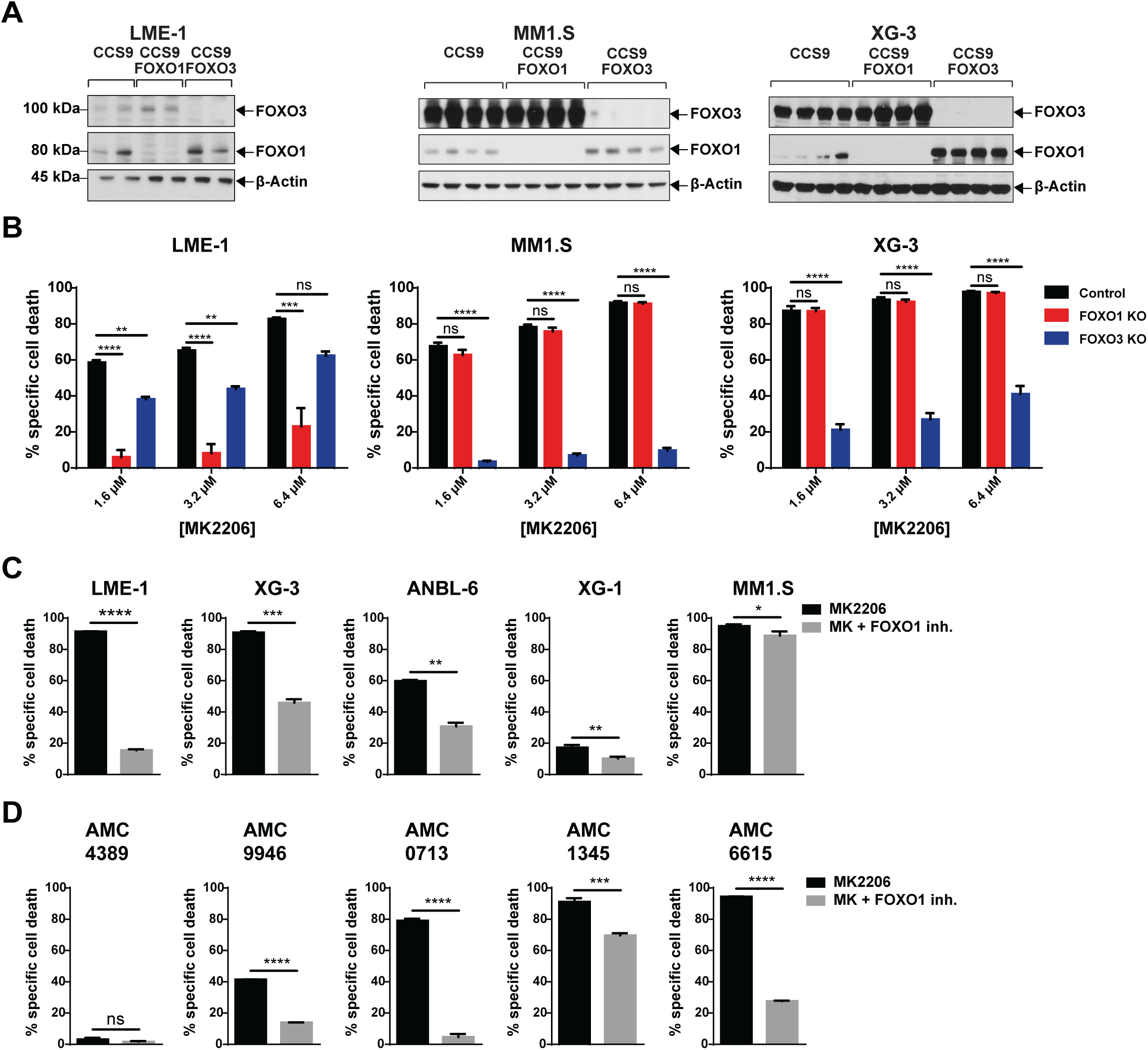
Cell death induced by AKT inhibition is dependent on FOXO1 or FOXO3 in MM cells. **(A)** Immunoblot analysis of CRISPR-Cas9 generated FOXO1 and FOXO3 knockout clones of the LME-1 (n=2), MM1.S (n=4), XG-3 (n=4) HMCLs. β-actin was used as loading control. **(B)** AKT inhibitor-induced cell death is dependent on the presence of FOXO1 in LME-1, and on FOXO3 in MM1.S and XG-3. Cloned knockout and control HMCLs were treated for 3 days with various concentrations of the MK2206 AKT inhibitor. 2 to 4 independently established clones were analyzed per condition. Red bars depict FOXO1 knockout clones, blue bars depict FOXO3 knockout clones. Means ± SEM of 3 independent experiments are shown (****p<0.0001; **p<0.01; ns = not significant; one-way ANOVA with Dunnet’s multiple comparison test). **(C)** AKT inhibitor-induced cell death in HMCLs can be rescued by FOXO1 inhibition (n=5). HMCLs were treated for 3 days with 3.2 μM MK2206 AKT inhibitor, with (grey bars) or without (black bars) 100 nM of the FOXO1 inhibitor AS1842856. Means ± SEM of 3 independent experiments are shown (****p<0.0001; ***p<0.001; **p<0.01; *p<0.05; ns = not significant; unpaired t-test with Welch’s correction). **(D)** Cell death of primary MM patient plasma cells induced by AKT inhibitor MK2206 (2.5 μM) can be overcome by the FOXO1 inhibitor AS1842856 (n=5). Cells were treated for 3 days with 3.2 μM MK2206 AKT inhibitor, with (grey bars) or without (black bars) 100 nM of the FOXO1 inhibitor AS1842856. Means ± SEM of 3 technical replicates are shown (****p<0.0001; ***p<0.001; ns = not significant; unpaired t-test with Welch’s correction). Specific cell death in these experiments was determined by 7-AAD viability dye staining and flow-cytometry.

We confirmed the tumor suppressive function of FOXO in MM using AS1842856, a small molecule inhibitor that blocks the transcriptional activity of FOXO1 and to a far lesser extent that of FOXO3^39^. In line with results observed in the FOXO-deficient cells, AS1842856 rescued LME-1 cells after MK2206 treatment but had almost no effect on the induced cell death in MM1.S cells. In other HMCLs, AS1842856 varyingly rescued MK2206-induced cell death **(Fig 2C)**. These results were confirmed in MM patient samples, where AS1842856 significantly inhibited MK2206-induced cell death in 4 out of 5 patient samples tested **(Fig 2D)**. The varying degree of rescue from cell death by AS1842856 may reflect the differential dependency on FOXO1 versus FOXO3 in these patients. Similarly, AS1842856 had no additional effect on FOXO1-deficient LME-1 cells, whereas AKT inhibitor-induced cell death of MM1.S cells, which required FOXO3, was partially rescued by AS1842856 **(Suppl Fig 2B)**. These results clearly demonstrate that FOXO transcription factors are crucial effectors of cell death upon inhibition of AKT, and function as AKT-restrained tumor suppressors in MM.

### Inhibition of AKT activates FOXO-controlled transcriptional regulation in MM cells

To determine FOXO-dependent transcriptional changes we performed gene expression profiling (GEP) of two independently established FOXO1-knockout clones from the LME-1 HMCL, and of two FOXO3-knockout clones from the MM1.S and XG-3 HMCLs, respectively. Cloned wildtype “Cas9 only” cells (WT) were used as controls. We anticipated that the three HMCLs would show considerable variance in the expression of FOXO target genes, due to the heterogeneous genetic backgrounds of these HMCLs. However, gene set enrichment analysis (GSEA) on the combined GEP datasets clearly indicated significant enrichment of a FOXO3 target gene set in the AKT inhibitor-treated WT control clones (‘WTMK’; <u>WT</u> clones, <u>MK</u>2206-treated) versus the untreated control and FOXO-knockout clones, and AKT inhibitor-treated FOXO-knockout clones (‘REST’) **(Fig 3A)**.

**FIGURE 3.**
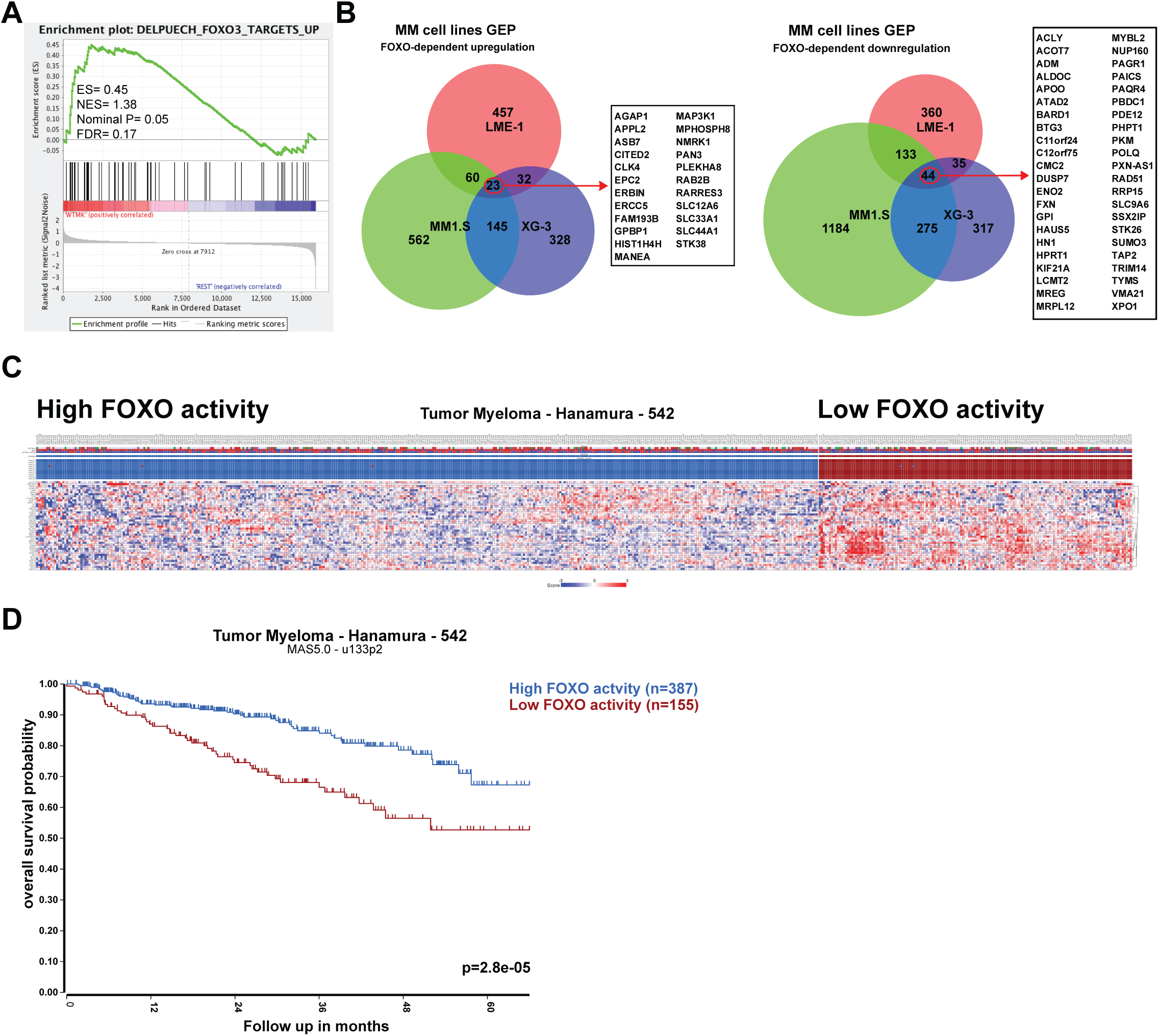
Inhibition of AKT induces a FOXO-dependent gene signature in MM cells. Independent LME-1 FOXO1 knockout clones (n=2), MM1.S FOXO3 knockout clones (n=2) and XG-3 FOXO3 knockout clones and their respective control clones (n=2) were treated overnight with 2.5 μM MK2206 AKT inhibitor, or left untreated, and subjected to gene expression profiling. **(A)** GSEA enrichment plot of upregulated FOXO3 target genes (DELPUECH_FOXO3_TARGETS_UP) in wildtype “cas9 only” (WT) clones treated overnight with 2.5 μM MK2206 (‘WTMK’; left side of the plot) versus MK2206-treated FOXO knockout clones, untreated WT and FOXO knockout clones (‘REST’; right side of the plot) from the LME-1, MM1.S and XG-3 HMCLs combined. False discovery rate (FDR), enrichment score (ES), normalized enrichment score (NES) and p-value are indicated in the enrichment plot. **(B)** Area proportional Venn diagrams depicting the number of genes that are upregulated (left panel) or downregulated (right panel) in a FOXO-dependent fashion upon AKT inhibition. Genes that overlap in all 3 HMCLs are listed alongside the Venn diagrams. Differentially expressed genes between the groups were defined based on p-values, using the following cutoffs: LME-1 p<0.15, MM1.S p<0.01, XG-3 p<0.02, (Annova corrected for multiple testing by false discovery rate). **(C)** K-means clustering results (10 rounds, 2 groups, blue and red boxes) and z-score heat maps based on the genes that are downregulated upon AKT inhibition in a FOXO-dependent fashion and overlapped in all 3 HMCLs (see **Fig 3B**, right) in a patient GEP dataset. This set contains gene expression profiling- and survival data of 542 MM patients. Blue depicts downregulated gene expression and red depicts upregulated gene expression. **(D)** Kaplan-Meier plot depicting overall survival of MM patients from the GEP dataset, using *k*-means clustering derived groups representing high and low expression of FOXO target genes (see **Fig 3B**).

Using variable cutoffs, we identified FOXO-regulated genes in MM1.S (848 up, 1541 down), XG-3 (329 up, 382 down) and LME-1 (438 up, 457 down) **(Suppl Table 1)**. In agreement, *k*-means unsupervised learning and principal component analysis (PCA) for MM1.S and XG3 showed that the MK2206-treated control clones (‘WTMK’) consistently clustered together versus a cluster consisting of the untreated control and FOXO3-knockout clones, and the MK2206-treated FOXO3-knockout clones (‘REST’). In contrast, the MK2206-treated control and FOXO1-deficient clones clustered together versus the untreated clones in LME-1. This may reflect intrinsic differences between LME-1 versus MM1.S and XG-3, and/or may indicate that FOXO1 has a less pronounced effect on gene expression compared to FOXO3 **(Suppl Fig 2A, B)**. Comparing these datasets, 23 genes were found to be consistently upregulated and 44 genes were downregulated upon AKT inhibition in a FOXO-dependent fashion **(Fig 3B)**, among which are the established direct FOXO targets *CITED2* (found in all three HMCLs) and *PIK3CA* (found in LME-1 and XG-3)^40,41^ **(Suppl Table 1)**. Despite this overlap, the majority of FOXO-dependent genes were specific for the different HMCLs, underscoring the heterogeneous and context-dependent nature of the transcriptional consequences of FOXO activation.

The FOXO-dependent downregulated genes found in all three HMCLs (FOXO_shared_down) were used to perform a *k*-means unsupervised learning analysis (2 groups, 10 rounds) on a MM patients GEP dataset (n=542) that includes clinical data^42^. Patients were clustered in 2 groups with respectively, low expression of FOXO suppressed genes (signifying high FOXO activity) versus high expression of FOXO suppressed genes (low FOXO activity) **(Fig 3C)**. Importantly, patients with high expression of FOXO suppressed genes, reflecting high AKT activity, show an inferior overall survival (p=0.000028) **(Fig 3D)**. The 90% survival was 9 months in this group with low FOXO activity versus 25 months in the high FOXO activity group, and the 2-year survival was 75% versus 91%. These results show that the loss of FOXO activity (and/or increased AKT activity) results in a more aggressive disease course, consistent with a tumor suppressive role of FOXO in MM.

### Inhibition of AKT induces a FOXO-dependent cell cycle arrest in HMCLs

Further inspection of the GSEA data on the combined datasets showed significant depletion of cell cycle and DNA replication/repair-associated gene sets in the MK2206-treated control clones **(Fig 4A, Supplemental Fig 4A)**, indicating that activation of FOXO induced cell cycle exit. In agreement, MM patients clustered according to high FOXO activity showed a significant depletion of these gene sets **(Suppl Fig 4B)**.

**FIGURE 4.**
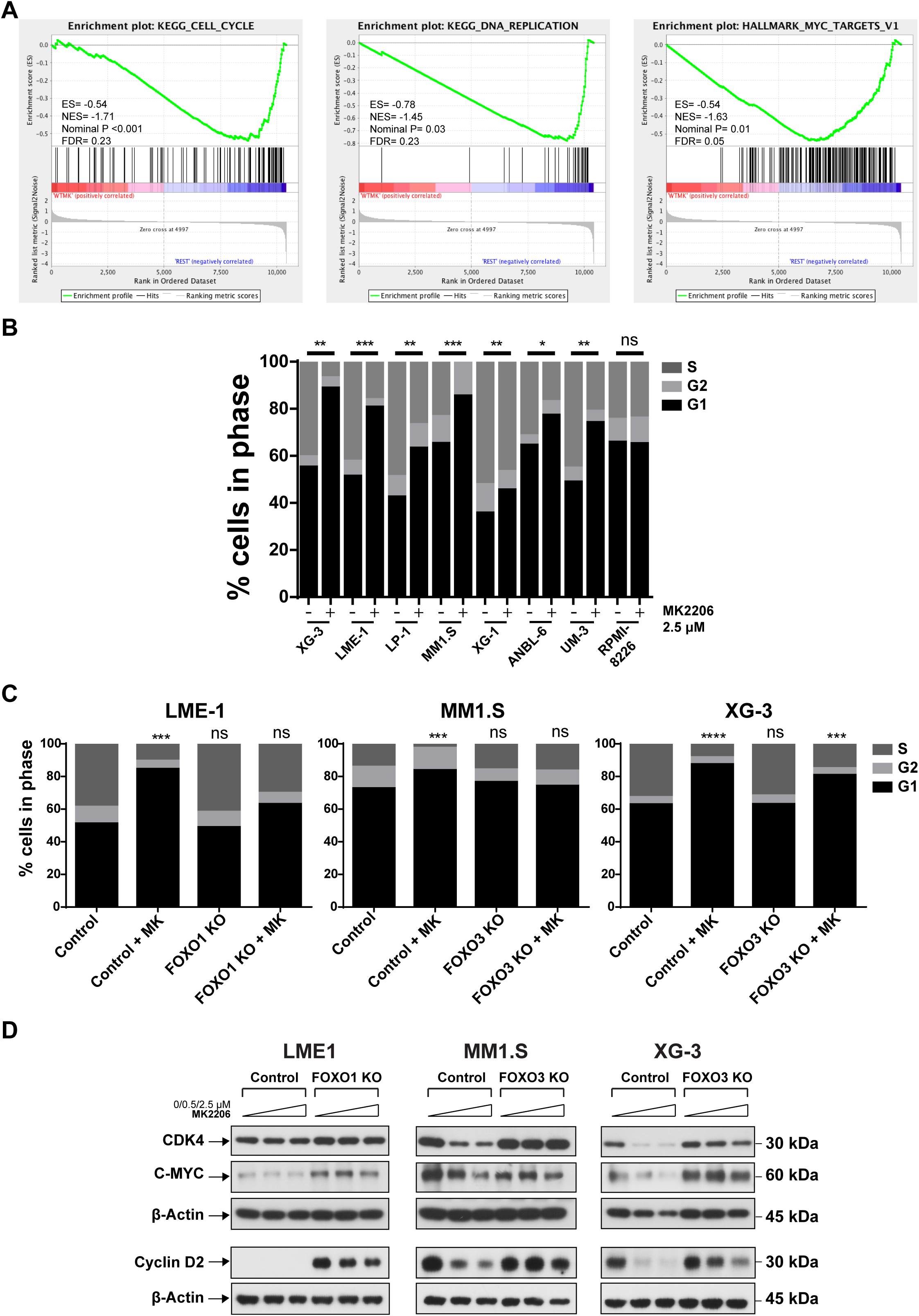
AKT inhibition impairs proliferation in a FOXO-dependent fashion in MM cells. **(A)** GSEA enrichment plots show a significant depletion of cell cycle and proliferation associated gene sets in wildtype “cas9 only” (WT) clones treated overnight with 2.5 μM MK2206 (‘WTMK’; left side of the plots) versus MK2206-treated FOXO knockout clones and untreated WT and FOXO knockout clones (‘REST’; right side of the plots). For GSEA, GEP datasets from the LME-1, MM1.S and XG-3 HMCLs were combined. False discovery rate (FDR), enrichment score (ES), normalized enrichment score (NES) and p-values are indicated in the enrichment plots. **(B)** BrdU incorporation cell cycle analysis of HMCLs (n=8) treated overnight with 2.5 μM MK2206. BrdU incorporation and DNA content was assessed by flow-cytometry. Sub G1 phase (dead) cells were excluded from the analysis. Percentages of cells in the G1, S, and G2 phase of the cell cycle are depicted. Statistical analysis (one-way ANOVA with Fisher’s Least Significant Difference post-test) was performed on the percentages of cells in S phase (***p<0.001; **p<0.01; *p<0.05; ns = not significant). The mean values of three experiments are depicted. **(C)** AKT inhibition leads to a FOXO-dependent G1 phase arrest. Cell cycle analysis of LME-1, MM1.S and XG-3 HMCLs and their respective FOXO1 or FOXO3 knockout clones treated overnight with 2.5 μM MK2206 (MK). Percentages of cells in the G1, S, and G2 phase of the cell cycle are depicted. Statistical analysis (one-way ANOVA with Bonferroni’s multiple comparison test) was performed on the percentages of cells in S phase compared to untreated control clones (****p<0.0001; ***p<0.001; ns = not significant). The mean values of 3 experiments are depicted. **(D)** Immunoblot analysis of cell cycle and proliferation associated proteins in LME-1, MM1.S and XG-3 control clones and their respective FOXO1 or FOXO3 knockout clones treated overnight with increasing concentrations of 0/0.5/2.5 μM MK2206. β-actin was used as loading control.

Correspondingly, cell cycle analysis showed that MK2206 treatment resulted in a significant loss of S phase and a concomitant increase in G1 phase in 7 out of 8 HMCLs tested **(Fig 4B)**, including the UM-3 HMCL that was unresponsive to AKT inhibition regarding cell death **(Fig 1A)**. Cell cycle exit was dependent on FOXO1 in LME-1 and on FOXO3 in MM1.S and XG-3 **(Fig 4C)**. Expression of the cyclin-dependent kinase 4 (CDK4) protein, which regulates G1 phase progression, was diminished by AKT inhibition in a dose-dependent manner in MM1.S and XG-3 control cells but not in FOXO3-deficient cells. In contrast, CDK4 remained largely unaffected in LME-1 cells. Protein expression of C-MYC was reduced in all three cell lines in a FOXO-dependent manner, which is also reflected in the GSEA analysis of the MYC targets geneset **(Fig 4A)**. Furthermore, C-MYC displayed higher basal protein levels in the FOXO1-deficient LME-1 cells and FOXO3-deficient XG-3 cells compared to control cells. Protein expression of Cyclin D2 was down modulated after AKT inhibition in a FOXO3-dependent manner in MM1.S and XG-3, but appeared to be activated in the FOXO1-deficient LME-1 cells **(Fig 4D)**. Analysis of GEP data indicated that the levels of *CDK4* mRNA were consistently down modulated by FOXO3-activation in MM1.S and XG-3, but not in LME-1 cells, suggesting that *CDK4* gene transcription is suppressed by FOXO3, but not FOXO1. In contrast, *C-MYC* mRNA levels were not affected in any of the HMCLs, whereas *CCND2* mRNA levels showed a FOXO3-dependent decrease in expression after AKT inhibition in XG-3 and MM1.S **(Suppl Fig 5)**. These data indicate that the effects of FOXO activation on the cell cycle involves both transcriptional and post-transcriptional mechanisms, which are context-dependent, but nonetheless result in a uniform cell cycle exit.

**FIGURE 5.**
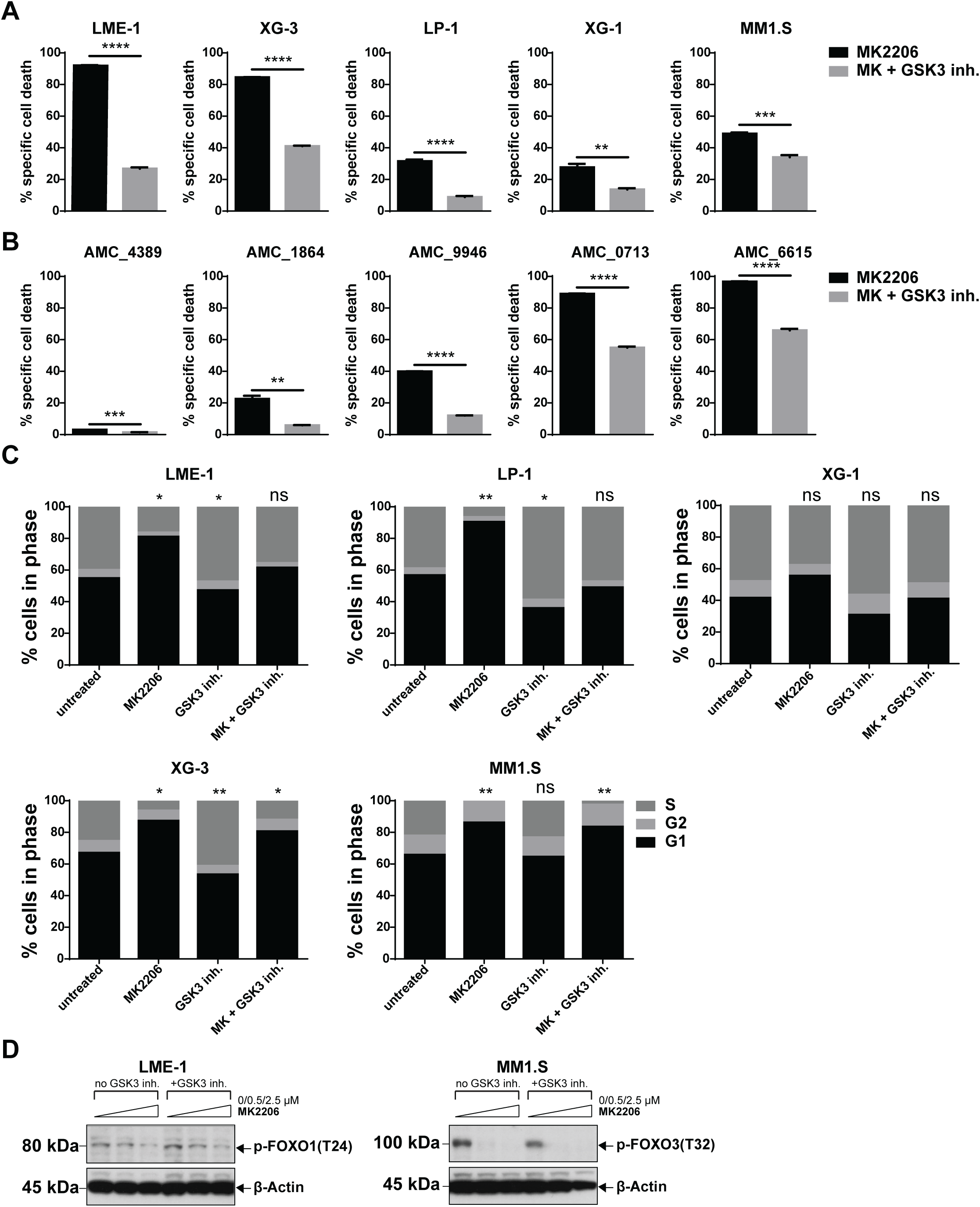
Inhibition of GSK3 partially rescues MM cells from AKT inhibitor-induced cell death and cell cycle arrest. **(A)** GSK3 inhibition partially rescued AKT inhibitor-induced cell death in HMCLs (n=5). Various concentrations of MK2206 were used for the different HMCLs (LME-1, LP-1 and XG-1: 3.2 μM; XG-3: 0.4 μM; MM1.S: 0.8 μM). Cells were co-treated with 1 μM CHIR99021 (GSK3 inh.) for 3 days. Means ± SEM of 3 independent experiments are shown (****p<0.0001; ***p<0.001; **p<0.01; unpaired t-test with Welch’s correction). **(B)** Partial rescue of AKT inhibitor-induced cell death in primary MM patient plasma cells (n=5). Cells were treated for 3 days with 2.5 μM MK2206 and 1 μM CHIR99021. Specific cell death was determined by 7-AAD viability staining and flow-cytometry. Means ± SEM of 3 technical replicates are shown (****p<0.0001; ***p<0.001; **p<0.01; unpaired t-test with Welch’s correction). **(C)** BrdU incorporation cell cycle analysis of HMCLs (n=5) treated overnight with 2.5 μM MK2206 and 1 μM CHIR99021. BrdU incorporation and DNA content was assessed by flow-cytometry. Sub G1 phase (dead) cells were excluded from the analysis. Percentages of cells in the G1, S, and G2 phase of the cell cycle are depicted. Statistical analysis (one-way ANOVA with Bonferroni’s multiple comparison test) was performed on the percentages of cells in S phase compared to untreated control cultures. Cultures were performed in triplicate (**p<0.01; *p<0.05; ns = not significant). **(D)** Immunoblot analysis of phospho-Thr24 FOXO1/phospho-Thr32 FOXO3 in LME-1 cells and MM1.S cells treated overnight with 0, 0.5 or 2.5 μM MK2206, with or without 1 μM CHIR99021. β-actin was used as loading control.

### GSK3 kinase activity is involved in AKT inhibitor-induced cell death and cell cycle arrest

GSK3 is an important physiological target of AKT that is inhibited by phosphorylation **(Fig 1C, D)**^20^. Activation of GSK3 downstream of AKT and PI3K inhibition has been implicated in apoptosis and cell cycle arrest^21,43^. Correspondingly, the GSK3-specific kinase inhibitor CHIR99021 significantly diminished cell death of AKT inhibitor-treated MM cells. Inhibition of GSK3 resulted in a partial rescue of MK2206-induced cell death, ranging from a 1.4-fold decrease in MM1.S cells, to a 3.4-fold decrease in LME-1 cells **(Fig 5A)**. A similar range of decrease in AKT inhibitor-induced cell death was observed in primary MM patient plasma cells co-treated with CHIR99021 **(Fig 5B).** GSK3 inhibition prevented AKT inhibitor-induced cell cycle arrest in the LP-1 and LME-1 HMCLs, whereas it had a modest effect in XG-3 and MM1.S. The effects of AKT inhibitor and GSK3 inhibitor treatment in XG-1 displayed a similar trend on the cell cycle as LME-1 and LP-1 but did not reach significance **(Fig 5C)**. Of note, in LME-1, LP-1 and XG-3 cells we observed a significant increase in S phase upon treatment with the GSK3 inhibitor alone, suggesting that constitutive AKT signaling in these HMCLs does not completely impede GSK3 kinase activity. The activation of FOXO1 or FOXO3 appeared not to be abrogated by GSK3 inhibition **(Fig 5D)**. These results indicate that GSK3 significantly contributed to the antimyeloma properties of AKT inhibition, acting in a cooperative fashion with FOXO transcription factors.

### GSK3 and FOXO activation upon AKT inhibition results in decreased MCL1 expression and sensitizes HMCLs for a selective MCL1 BH3 mimetic

Previously, it was shown that AKT and GSK3 kinase activity are involved in apoptosis by regulating the protein stability of MCL1^26,44,45^, a BCL2-family member that is essential for the survival of non-malignant plasma cells and myeloma cells^27,46^. These findings prompted us to investigate the effects of AKT inhibition on MCL1 protein levels in MM. MCL1 protein expression was diminished by AKT inhibitor treatment in responsive HMCLs (LME-1, MM1.S and XG-3), which depended on FOXO and GSK3 activity **(Fig 6A)**. Similarly, MCL1 protein expression was decreased in AKT inhibitor-treated primary MM patient plasma cells **(Fig 6B)**. Cycloheximide chase experiments showed that AKT inhibitor treatment increased MCL1 protein turnover in responsive HMCLs **(Fig 6C)**. In contrast, BCL2 and BCL-XL protein stability remained unchanged after AKT inhibition in all tested HMCLs **(Suppl Fig 6A)**. In agreement, overexpression of MCL1 in LME-1, MM1.S and XG-3 prevented AKT inhibitor-induced cell death **(Fig 6D, E)**. The S63845 small molecule inhibits MCL1 and displays potent antimyeloma acitivity^47^. Based on our results we asked whether AKT inhibition sensitized myeloma cells for this MCL1 inhibitor. We exposed the MK2206-responsive HMCLs LME-1, MM1.S, XG-3 and the unresponsive HMCLs UM-3 and RPMI-8226 to increasing concentrations of MK2206, S63845 and the combination of both. A clear potentiating effect on induced cell death was observed with the combination of inhibitors, even in the MK2206 unresponsive HMCLs **(Fig 6F, Suppl Fig 6B)**.

**FIGURE 6.**
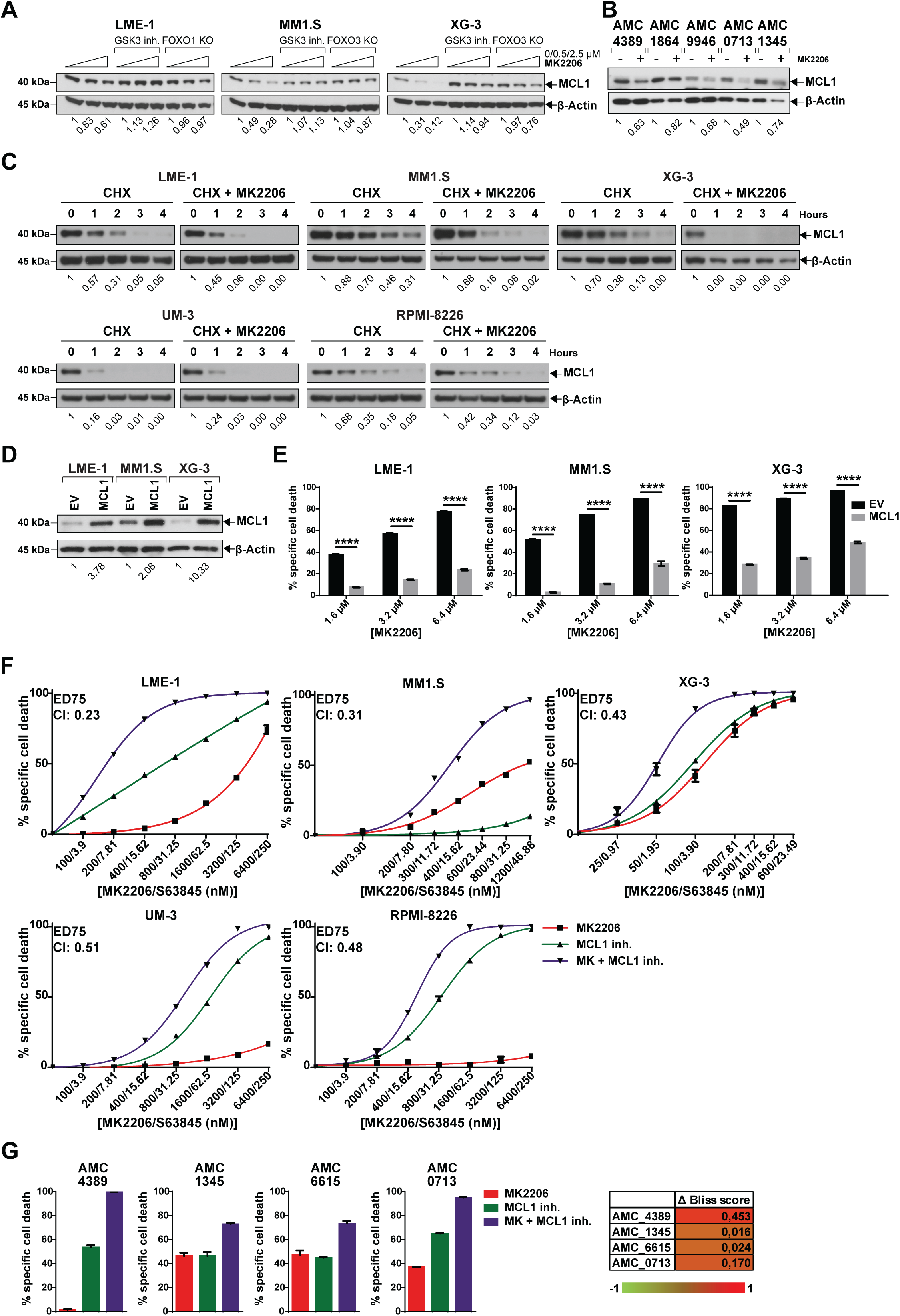
AKT inhibition in MM cells leads to the FOXO/GSK3-mediated MCL1 down modulation resulting in cell death. **(A)** Immunoblot analysis of MCL1 protein expression in wildtype “cas9 only” clones treated overnight with increasing concentrations of MK2206 (0; 0.5; 2.5 μM) with or without 1 μM CHIR99021, and in MK2206-treated (0; 0.5; 2.5 μM) FOXO1 or FOXO3 knockout HMCLs. Immunoblot analysis of MCL1 protein expression in primary MM patient plasma cells (n=5) treated overnight with 2.5 μM MK2206. **(C)** Immunoblot analysis of MCL1 protein stability in HMCLs (n=5) after cycloheximide (CHX) treatment (200 µg/ml), with or without pretreatment of 2.5 µM MK2206 for 12 hours. Cells were treated with CHX as indicated by depicted time points. **(D)** Immunoblot analysis of MCL1 protein expression in HMCLs (n=3) overexpressing MCL1. HMCLs expressing empty vector were used as controls. β-actin was used as loading control. **(E)** MCL1 overexpression rescues AKT inhibitor-induced cell death. HMCLs overexpressing MCL1 (n=3) were cultured for 3 days with various concentrations of MK2206 (1.6; 3.2; 6.4 µM). HMCLs transduced with empty vector were used as controls. Specific cell death was assessed by 7-AAD viability dye staining and flow-cytometry. Means ± SEM of 3 independent experiments are shown (****p<0.0001; one-way ANOVA with Bonferroni’s multiple comparison test). **(F)** Inhibition of AKT sensitizes HMCLs (n=5) to MCL1 inhibitor induced cell death. Cells were treated for 3 days with various concentrations of MK2206 and S63845 (MCL1 inh.), as indicated on the x-axis of the graphs. Percentages of specific cell death are depicted. Means ± SEM of 3 experiments are shown. Chou-Talalay combination index (CI) values at ED75 (effective dose causing 75% cell death) are indicated in the graphs. **(G)** Inhibition of AKT potentiates for MCL1 inhibitor-induced cell death in primary patient plasma cells (n=4). Cells were treated for 3 days with a concentration of MK2206 and S63845 (2,5 µM MK2206, 100 nM S63845 for AMC_4389, 100nM MK2206, 4nM S63845 for AMC_1345 and AMC_6615, 500 nM MK2206, 20nM S63845 for AMC_0713) either as single drug or a combination. Means ± SEM of three technical replicates are shown. The **Δ** Bliss score was calculated by subtracting the predicted cell death (Bliss) from the actual observed effect of the combined inhibitors, −1 indicates an antagonistic effect and +1 indicates a synergistic effect.

In accordance with the observed results in HMCLs, the combination of AKT and MCL1 inhibitors resulted in cell death consistently higher than the predicted Bliss score in primary MM cells, indicating a potentiating effect of this drug combination **(Fig 6G)**. These results indicate that the FOXO- and GSK3-mediated decrease in MCL1 protein expression after AKT inhibition sensitizes myeloma cells for the MCL1-specific inhibitor S63845, improving the efficacy of these novel therapeutic modalities.

## DISCUSSION

Here we showed that the FOXO1 and FOXO3 transcription factors and the GSK3 kinase act as AKT-repressed tumor suppressors in MM cells. As such, FOXO1/3 and GSK3 are critical mediators of the antimyeloma effects of AKT-targeted therapy. In agreement, we showed that the expression levels of a set of FOXO targets genes is related to overall survival rates in MM patients; low FOXO activity (reflecting high AKT activity) identifies a patient subgroup with inferior survival.

We demonstrated that upon AKT inhibition FOXO1/3 and GSK3 mediate cell cycle exit by repressing genes involved in DNA replication and cell cycle progression, and cause cell death by provoking the loss of MCL1 protein expression. These observations offer important leads to improve therapeutic strategies aimed at the PI3K/AKT pathway. As an example, we showed that the AKT inhibitor MK2206 synergized with the recently developed MCL1 inhibitor S63845^47^. Combination of these two drugs very efficiently caused cell death of MM cells, even in cells refractory to AKT inhibition, warranting further investigation into the clinical efficacy of such combination therapies. Targeting AKT and MCL1 simultaneously can be considered a vertical inhibition strategy, in which two points of the same pathway are inhibited. Similar vertical inhibition strategies for the PI3K/AKT/mTOR pathway have shown to be synergistic in multiple cancer types^48–50^.

The tumor suppressive roles of FOXO1/3 and GSK3 partly explain the constitutive activation of the PI3K/AKT pathway in MM cells, underscoring its crucial function in tumor cell survival. Whether PI3K/AKT signaling has a similar role in the maintenance of normal plasma cells remains unknown. However, AKT activity was shown to be important in the development of normal plasma cells, as *in vitro* differentiation of mouse plasma cells was inhibited by forced expression of constitutive active FOXO1. Conversely, FOXO1 knockout in mouse mature B cells or treatment with PI3K inhibitors increased plasma cell formation^12,51^. In contrast, expression of constitutive active FOXO1 in classical Hodgkin lymphoma directly drove *PRDM1/BLIMP1* expression, thereby activating a plasma cell gene signature^52^. The role of FOXO3 in plasma cell differentiation is less clear, whereas FOXO3-deficient mouse mature B cells show no apparent defects in plasma cell development^13^. However, expression of FOXO3 increases from germinal center B cells to plasma cells^53^. These observations suggest that FOXO transcription factors act in a context-dependent fashion in normal and malignant plasma cells. This is emphasized by our GEP data, showing relatively limited overlap between FOXO-regulated genes in three different HMCLs. Despite this apparent heterogeneity, the effects of FOXO activation on proliferation and cell death were remarkably uniform, as reflected by gene expression enrichment analysis performed on the combined datasets. There are some reports that suggest that FOXO1 and FOXO3 are functionally linked and act redundantly, for instance in autophagy^54^ and development of thymic lymphomas and hemangiomas^55^, whereas in lymphocyte development these FOXO transcription factors have specialized as well as redundant functions^12,13^. However, in MM cells the functions of FOXO1 and FOXO3 suggest there is no obvious overlap, since the loss of FOXO1 was not compensated by FOXO3, or vice versa, despite increased expression of the alternate family member.

A major difference between normal and malignant plasma cells is their proliferative capacity, which can be attributed to recurrent genomic abnormalities that result in the aberrant expression of cell cycle related genes, such as D-type cyclins and C-MYC, which are nearly universal events in MM^56^. Despite aberrant expression, these cell cycle-associated genes were nevertheless repressed by FOXO1/3 upon AKT inhibition, thereby reversing the oncogenic proliferative program of MM cells. Pharmacological inhibition of AKT resulted in a FOXO-dependent G1 phase arrest in MM cells, consistent with earlier reports on lymphomas and other cancer cell types^57–60^. As previously shown, FOXO may cause cell cycle arrest by driving the expression of the cyclin-dependent kinase inhibitor p27(kip1)^61^, and by reducing the expression of D-type cyclins^62^. We found that AKT inhibition resulted in a FOXO-dependent down modulation of cyclin D2, CDK4 and C-MYC protein expression, whereas p27(kip1) was not affected (data not shown). In agreement, GSEA indicated that MYC target genes were significantly depleted from AKT inhibitor-treated control cells **(Suppl Fig 4)**. In addition, DNA repair gene expression signatures were significantly down modulated upon activation of FOXO1/3 in MM **(Suppl Fig 4)**, suggesting that combining AKT inhibitors with DNA-damaging agents might be a promising treatment option for MM patients.

Our date emphasizes the pivotal importance of FOXO transcription factors and GSK3 kinase activation on MM cell survival and cell cycle progression downstream of AKT inhibition. AKT-mediated phosphorylation of FOXO1 or FOXO3 was not affected by GSK3 kinase inhibition, suggesting that GSK3 did not act upstream of FOXO1/3. These data indicate that FOXO1/3 and GSK3 act in a cooperative fashion. The role of GSK3 in the regulation of cell death downstream of PI3K/AKT signaling was described previously in various types of cancer^21,63–66^. However, the role of GSK3 in MM is less clear, as both prosurvival and proapoptotic functions have been ascribed to this kinase^67–73^. To our knowledge, our results are the first to indicate that GSK3 is an important mediator of cell death in MM cells controlled by AKT signaling. The partial and heterogeneous effect of GSK3 inhibition most likely reflects the molecular heterogeneity of the HMCLs and patient samples used in this study.

Recently, it has become clear that MM displays marked clonal heterogeneity, and that the tumor consist of subclonal variants that undergo clonal evolution during treatment and progression. Moreover, the mutation spectrum also may change over time, alluding to ongoing mutagenic processes that affect new candidate genes involved in therapy resistance and disease progression^74,75^. Based on our data, it is conceivable that treatment aimed at the PI3K/AKT pathway in MM may result in the selection, or appearance, of subclonal variants that harbor mutations inactivating FOXO and/or GSK3. Currently, several clinical trials assessing the efficacy of AKT inhibitors for the treatment of MM are underway. Our data underscores that inhibition of AKT can induce tumor cell vulnerabilities that can be exploited therapeutically, such as the AKT mediated negative regulation of MCL1.

## ACKNOWLEDGEMENTS

This research was supported by the Netherlands Organization for Scientific Research Innovational Research Incentives Scheme VIDI grant no.16126355, and by grant AMC 2018-11597 of the Dutch Cancer Society (both to J.E.J.G)

## AUTHORSHIP AND CONFLICT-OF-INTEREST STATEMENTS

J.E.J.G. and T.A.B. designed the research; T.A.B., G.d.W., C.M. and J.E.J.G. performed the experiments; E.E., R.J.B., C.J.v.N., S.T.P., M.S. and J.E.J.G. analyzed the data; T.A.B., G.d.W. and J.E.J.G. wrote the manuscript; and all authors edited the manuscript.

E.E. received research funding from Hoffman-La Roche Ltd. and from Gilead Sciences Inc.

