## Supplementary material for "Targeting AKT elicits tumor suppressive functions of FOXO transcription factors and GSK3 kinase in Multiple Myeloma": Suppl methods and Figures

*Constructs and retroviral/lentiviral transductions*

To generate CRISPR/Cas9 knockout (KO) HMCL clones, the pL-CRISPR.EFS.GFP vector was used. The pL-CRISPR.EFS.GFP was a gift from Benjamin Ebert (Addgene plasmid #57818). Control clones were established using the empty pL-CRISPR.EFS.GFP vector. To generate FOXO1 knockouts, 3 different gRNAs were transduced simultaneously (targeting sequences were GCTCGTCCCGCCGCAACGCG, CCCGGCCGCGCTCGTGCACC and GTTGCCCCACGCGTTGCGGC). For FOXO3 knockouts, we used 2 different gRNAs (targeting sequences ACTGCATAGTCGATTCATGC and TGGCCGCCTGTCGCCCATCA). Lentiviral particles were produced by transfecting the lentiviral vector together with psPAX2 (Addgene plasmid #12260) and pMD2.G/VSVG (Addgene plasmid #12259), both gifts from Didier Trono, into HEK293T cells (ratio; 4:2:1), using Genius transfection reagent (Westburg, Leusden, the Netherlands) according to manufacturer's protocol. Viral supernatant with the addition of 8 µg/mL polybrene (Sigma Aldrich, St. Louis, MO) was spinfected onto the HMCLs and incubated overnight. Five days after transduction the cells were analyzed for GFP expression on a FACSCANTO II flow cytometer (BD Biosciences, Franklin Lakes, NJ) and subsequently, GFP positive cells were sorted in 96-wells plates at a density of 10 cells per well using a Sony SH800S cell sorter (Sony Biotechnology, San Jose, CA).

LZRS-*MCL-1*-IresGFP was created by subcloning *MCL-1* obtained from pCDNA3.1-*MCL-1* using BamHI/XbaI, into LZRS-IresGFP using the BamHI/XhoI sites. Retroviral particles were obtained by transfecting Phoenix-Galv cells with the retroviral vectors using Genius transfection reagent according to manufacturer's protocol. The viral supernatant was then spun down on 24-wells plates coated with Retronectin (Takara

Bio Inc, Kusatsu, Japan) according to manufacturer's protocol. The viral supernatant was removed and the HMCLs were incubated in the virus coated plates for 3 days in fresh IMDM +10% FCS. Cells were further expanded for 5 more days and then FACS sorted, sorting only the top 15% GFP expressing HMCLs.

*Immunoblotting*

Cells ($2 \times 10^6$) were washed with ice-cold PBS and lysed in ice-cold radioimmunoprecipitation assay (RIPA) lysis buffer (50 mM Tris-HCl pH 7.4, 150 mM of NaCl, 1% Nonidet P-40, 0.5% Na-Deoxycholate, 0.1 % Sodium Dodecyl Sulfate), supplemented with protease and phosphatase inhibitors (EDTA-free protease mixture inhibitor; Roche Diagnostics, Rotkreuz, Switzerland) (PhosSTOP, Roche Diagnostics). Cell lysates were homogenized by passing them through a 27G needle and protein concentrations were measured using the bicinchoninic acid assay (Sigma Aldrich). For each sample, 10-15 µg of protein lysate was used for protein separation in Precise 4–20% gradient Tris-SDS gels (Thermo Fisher Scientific, Waltham, MA). Separated protein lysates were transferred onto polyvinylidene difluoride membranes (Immobilon-P; EMD Millipore, Burlington, USA); membranes were blocked in tris-buffered saline (TBS) + 5% bovine serum albumin (BSA) and 0.1% Tween 20 (TBS-T) or in TBS-T + 5% milk for 1 H at room temperature (RT) Primary Antibodies were incubated overnight at 4°C. After washing in TBS-T, the membranes were incubated for 1 H at RT with secondary antibodies (goat anti-rabbit HRP or rabbit anti-mouse HRP; DAKO, Agilent Technologies, Heverlee, Belgium). Protein expression was visualized by use of Amersham ECL Prime Western blotting detection reagent (GE Healthcare Bio-Sciences AB, Uppsala, Sweden). The primary antibodies used in this study are: Phospho-FoxO1 (Thr24)/FoxO3a (Thr32), FoxO1 (C29H4), FoxO3a (75D8), Phospho-

Akt (Ser473) (D9E) XP, Phospho-Akt (Thr308) (D25E6) XP, Akt1/2/3 (C67E7), mTor (7C10), Phospho-mTor (Ser2448), Phospho-GSK-3β (Ser9) (D85E12) XP, GSK-3α/β (D75D3), Phospo-S6 (D57.2.2E), S6 (5G10), CDK4 (D9G3E), Cyclin D2 (D52F9), BCL2 (124), BCL-XL (54H6), (Cell signaling technologies, Beverly, MA), C-Myc (EP121) (Epitomics, Burlingame, CA), MCL-1 (Y37) (Abcam, Cambridge, UK), β-Actin (C4) (Milipore).

*Flow cytometry*

To determine specific cell death, cells were plated in 96-wells plates (5 x $10^4$ or 1 x $10^5$ cells per well) and treated as described in figure legends. After treatment, cells were stained with 7-aminoactinomycin-D (7-AAD; Biolegend Inc, San Diego, CA) for 10' at RT and measured by flow cytometry. CD38 expression was determined by staining cells for 30' with anti CD38-APC (HIT2) (BD Biosciences) and measured by flow cytometry. Debris and doublets were excluded based on forward and side scatter height versus width profiles. Cell death was based on the 7-AAD incorporation. To determine the percentages of early apoptosis, late apoptosis and necrosis, cells were stained with Annexin V-GFP (IQProducts, Groningen, the Netherlands) in Annexin binding buffer (10mM Hepes, 150 mM NaCl, 5 mM KCl, 1.8 mM $CaCl_2$, 2,1 mM $MgCl_2$, 1% glucose, 0.5% BSA) and To-Pro-3 (Molecular Probes, Eugene, OR). Data was analyzed using FlowJo (Tree Star Inc, Ashland, OR) and specific cell death was calculated as previously described[28].

To assess effects on the cell cycle of HMCLs, cells were plated at a density of 1 x $10^6$ cells/ml and cultured O/N. Subsequently the cells were incubated 1 H with 20 μM bromodeoxyuridine (BrdU; Sigma Aldrich), washed with ice cold PBS and fixed in ice cold 75% ethanol at least O/N. The fixed cells were then treated for 25' at 37°C with

0.4 mg/ml pepsin (Sigma Aldrich) dissolved in 0.2 mM HCl. Thereafter, the cells were incubated in 2 M HCl for 25’ at 37°C and washed once in 0.5% BSA in PBS (PBS-B) followed by another washing step with 0.5 % tween in PBS-B (PBS-TB) and stained using anti-BrdU FITC (clone B44; BD Bioscience) for 30’ at RT. After subsequent washing with TBS-B and TBS-TB the cells were incubated for 15’ at 37°C with 0.1 µM TO-PRO-3-Iodide (Invitrogen life technologies, Carlsbad, CA) and 500 µg RNAse A (Sigma Aldrich) dissolved in PBS supplemented with 0.02% $NaN_3$, 0.5% BSA and measured by flow cytometry.

### SUPPLEMENTAL FIGURE 1

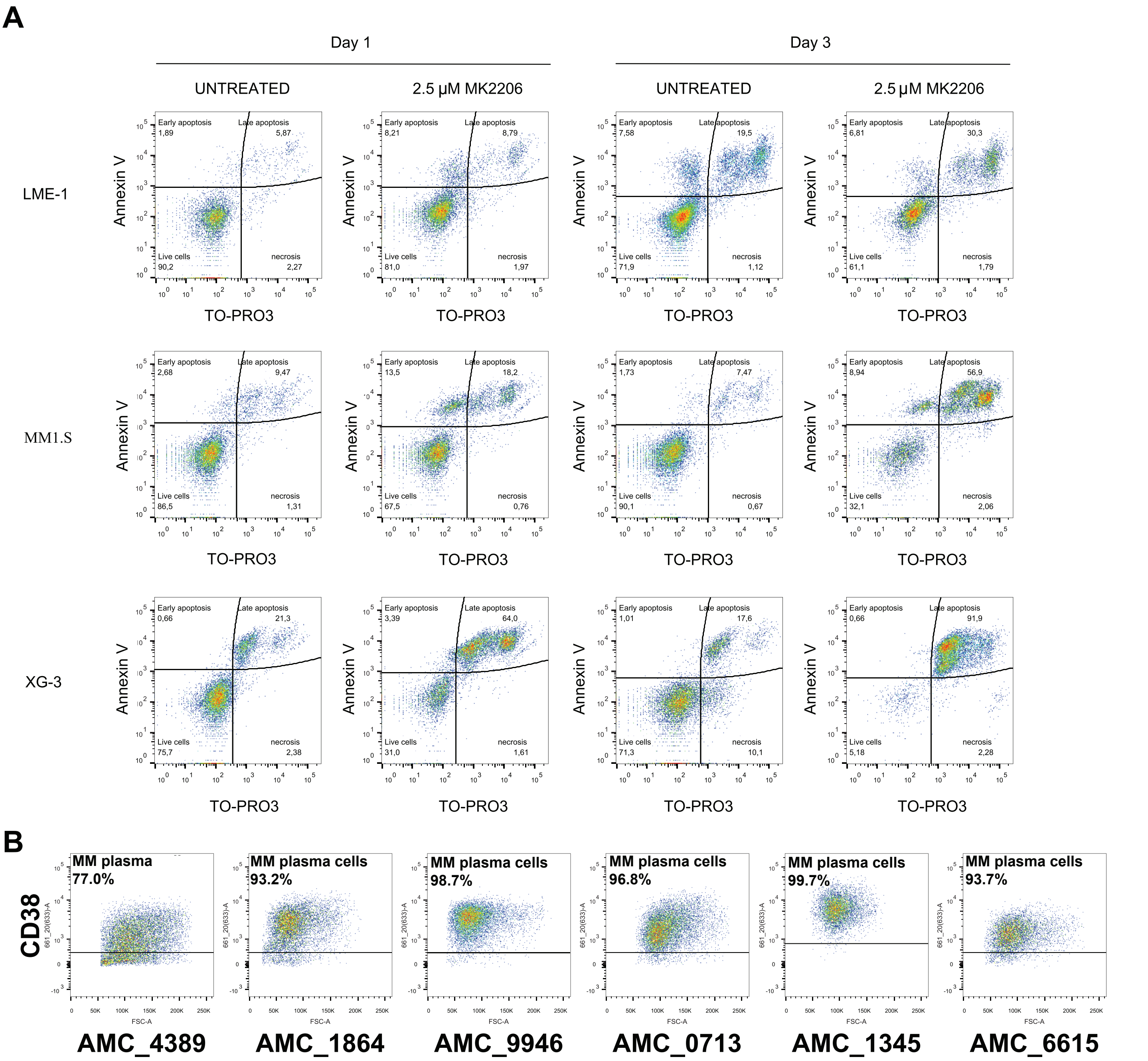


**SUPPLEMENTAL FIGURE 1. HMCLs Annexin V staining after MK2206 treatment, Primary MM patient plasma cell enrichment. (A)** HMCLs were treated for 1 and 3 days with 2.5 µM MK2206. Next the cells were stained with FITC-conjugated Annexin V and ToPro3 and analyzed by flow cytometry. Percentages of Annexin V single (early apoptosis), To-Pro-3 single (necrosis) and double positive cells (late apoptosis) are indicated in the FACs plots. **(B)** Primary MM plasma cells were enriched by Ficoll density centrifugation primary samples obtained from MM patients (n=6). After enrichment, samples were stained with APC-conjugated anti-CD38 and analyzed by flow cytometry. Percentages of CD38 positive cells are shown in the FACS plots.

### SUPPLEMENTAL FIGURE 2

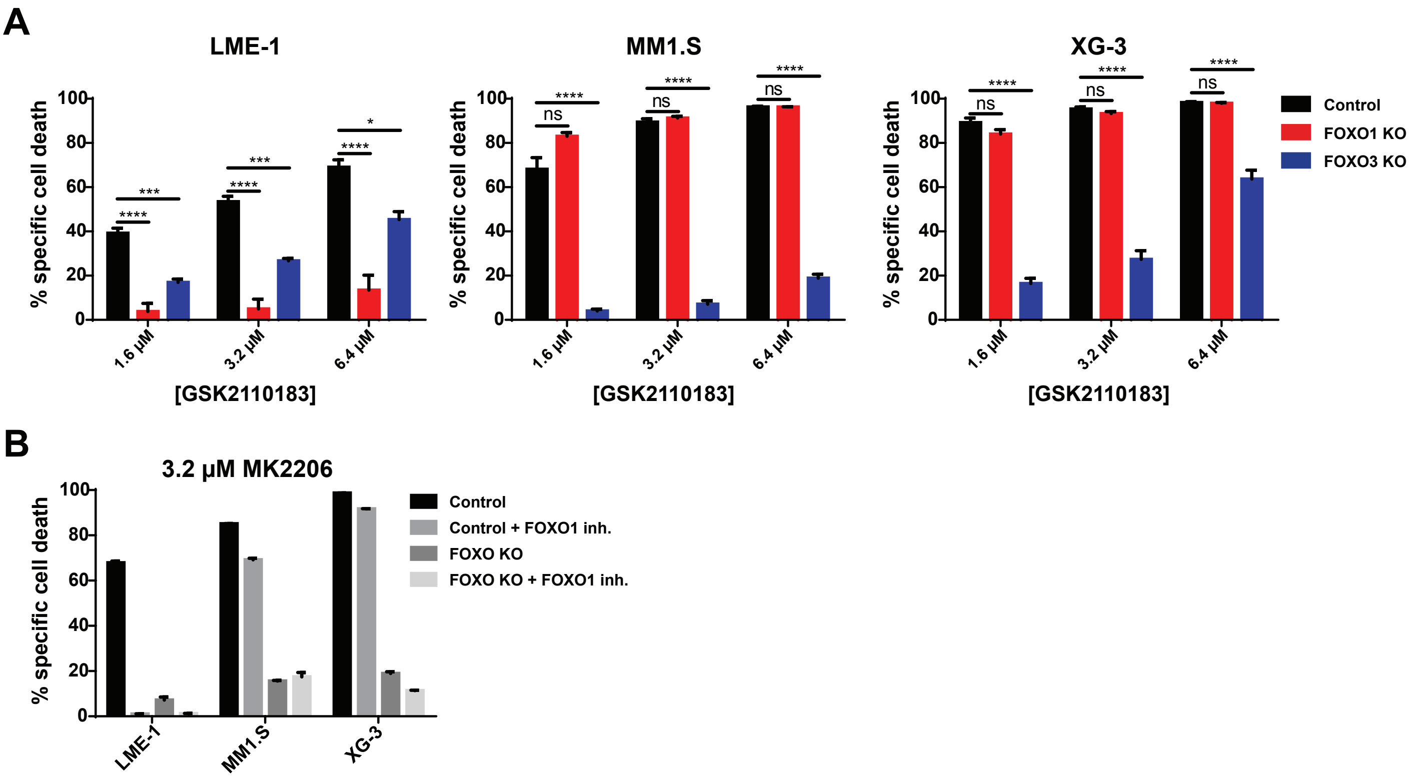


**SUPPLEMENTAL FIGURE 2. Cell death induced by GKS2110183 in HCMLs is also dependent on FOXO transcription factors, specificity assessment of FOXO1-inhibitor AS1842856**

**(A)** GSK2110183 induced cell death is dependent on the presence of FOXO1 in LME-1, and on FOXO3 in MM1.S and XG-3. Cloned knockout and control HMCLs were treated for 3 days with various concentrations of the GSK2110183 AKT inhibitor. 2 to 4 independently established clones were analyzed per condition. Red bars depict FOXO1 knockout clones, Blue bars depict FOXO3 knockout clones. Means ± SEM of 3 independent experiments are shown (****p<0.0001; ***p<0.001 **p<0.01; *p<0.05, ns = not significant; one-way ANOVA with Dunnet's multiple comparison test). **(B)** AS1842856 rescued FOXO1-dependent cell death in LME-1 but had no effect on FOXO3-dependent cell death in MM1.S and XG-3 cells. FOXO1 KO cells (LME-1), FOXO3 KO cells (MM1.S) and their respective control clones were treated with 2.5 µM MK2206 for 3 days with or without 100 nM of the AS1842856 FOXO1 inhibitor. Specific cell death was determined by 7-AAD incorporation followed by flow cytometry. Means ± range of the technical replicates are shown

### SUPPLEMENTAL FIGURE 3

A

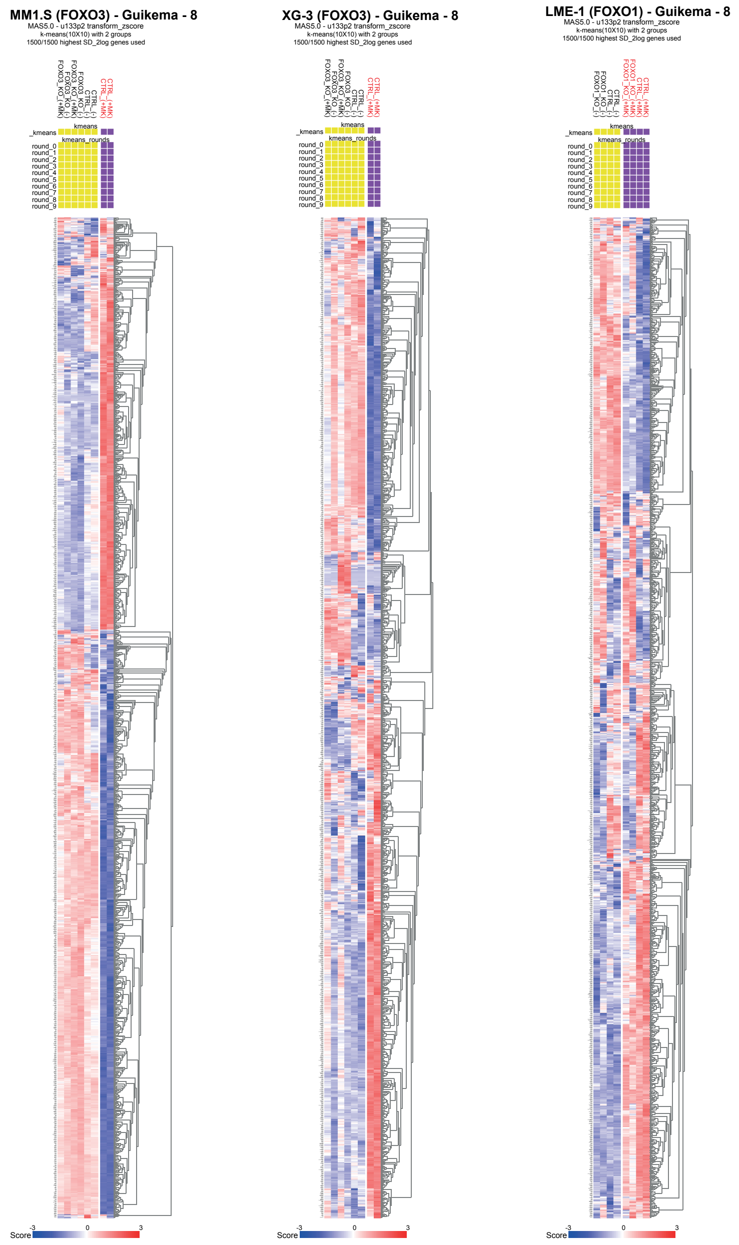


B

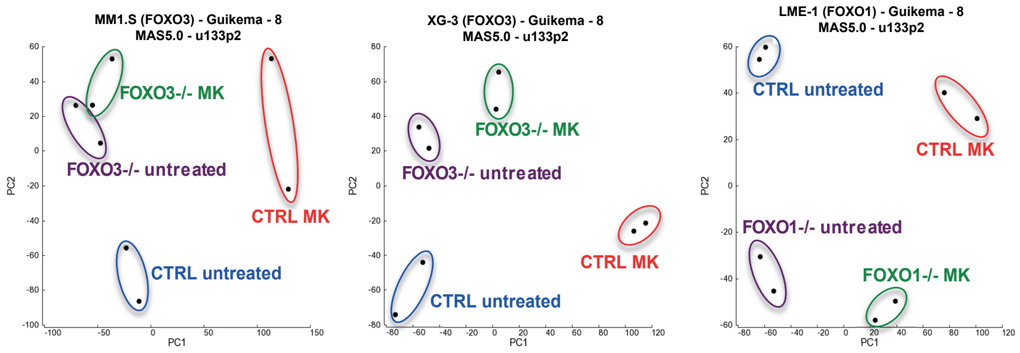


**SUPPLEMENTAL FIGURE 3. K-means clustering and principal component analysis of HMCLs**
Independently established LME-1 FOXO1 knockout clones (n=2), MM1.S FOXO3 knockout clones (n=2) and XG-3 FOXO3 knockout clones and their respective control clones (n=2) were treated overnight with 2.5 μM MK2206 AKT inhibitor, or left untreated, and subjected to gene expression profiling. A total of 8 samples was used for gene expression profiling: 2x untreated control clones: CTRL_(-); 2x MK2206-treated control clones: CTRL_(+MK); 2x untreated FOXO-knockout clones: FOXO_(-); 2x MK2206-treated FOXO-knockout clones: FOXO_(+MK). 10 rounds of K-means clustering to divide samples into 2 groups were performed on 1500 genes that showed the highest standard deviations (SD) in these 8 samples. **(A)** K-means clustering results (yellow and purple boxes) and z-score heat maps of the 1500 highest SD genes are shown. Blue depicts downregulated gene expression and red depicts upregulated gene expression. **(B)** Principle component analysis of z-score transformed expression data shows that AKT-inhibitor treatment and FOXO-status are the major determinants of variation in gene expression. Independent clones that were treated in a similar fashion are indicated in colored circles.

### SUPPLEMENTAL FIGURE 4

**A**

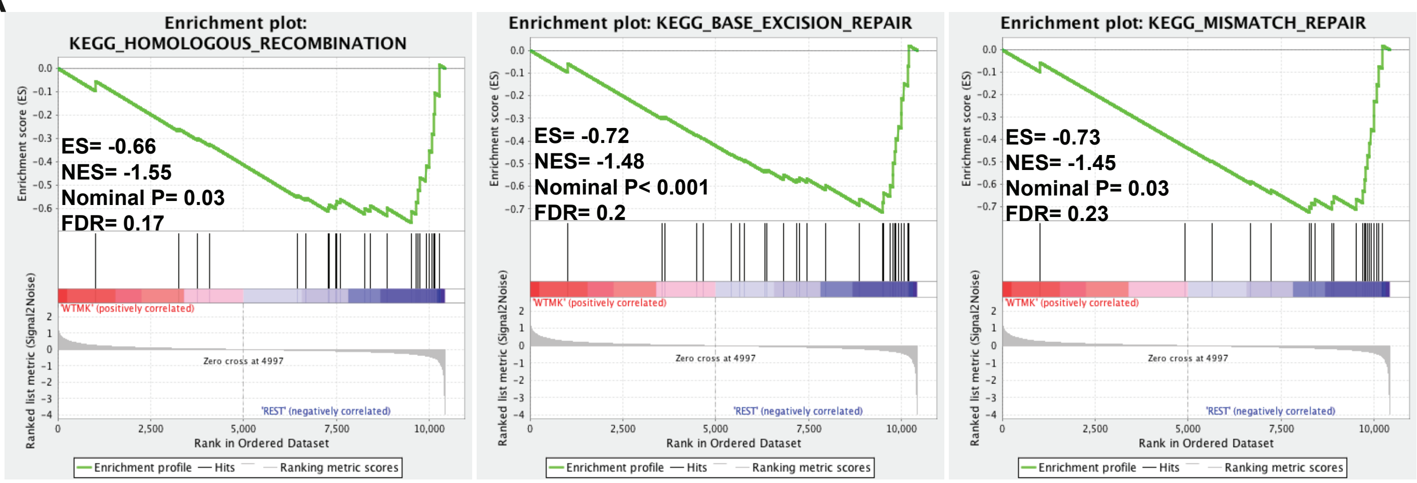


**B**

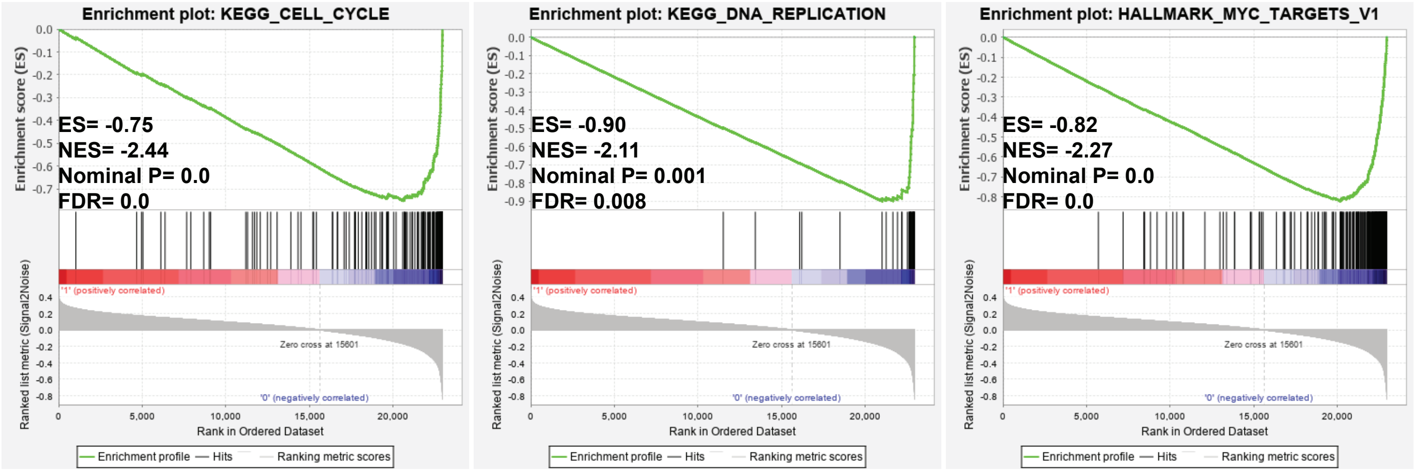


**SUPPLEMENTAL FIGURE 4. Gene set enrichment analysis shows that FOXO activation down modulates expression of DNA repair genes and C-MYC target genes**

**(A)** GSEA enrichment plots show a significant depletion of DNA repair associated gene sets and target genes in control (WT) clones treated overnight with 2.5 µM MK2206 ('WTMK'; left side of the plots) versus MK2206-treated FOXO knockout clones and untreated WT and FOXO knockout clones ('REST'; right side of the plots). For GSEA, GEP datasets from the LME-1, MM1.S and XG-3 HMCLs were combined. False discovery rate (FDR), enrichment score (ES), normalized enrichment score (NES) and p-value are indicated in the enrichment plots. **(B)** GSEA enrichment plots show a significant depletion of cell cycle associated gene sets and MYC target genes in patients clustered in the high FOXO activity ("1", left side of the plots) versus patients clustered in the low FOXO activity ('0'; right side of the plots). Groups were clustered using the FOXO_shared_down geneset as described in Figure 3B. False discovery rate (FDR), enrichment score (ES), normalized enrichment score (NES) and p-value are indicated in the enrichment plots.

### SUPPLEMENTAL FIGURE 5

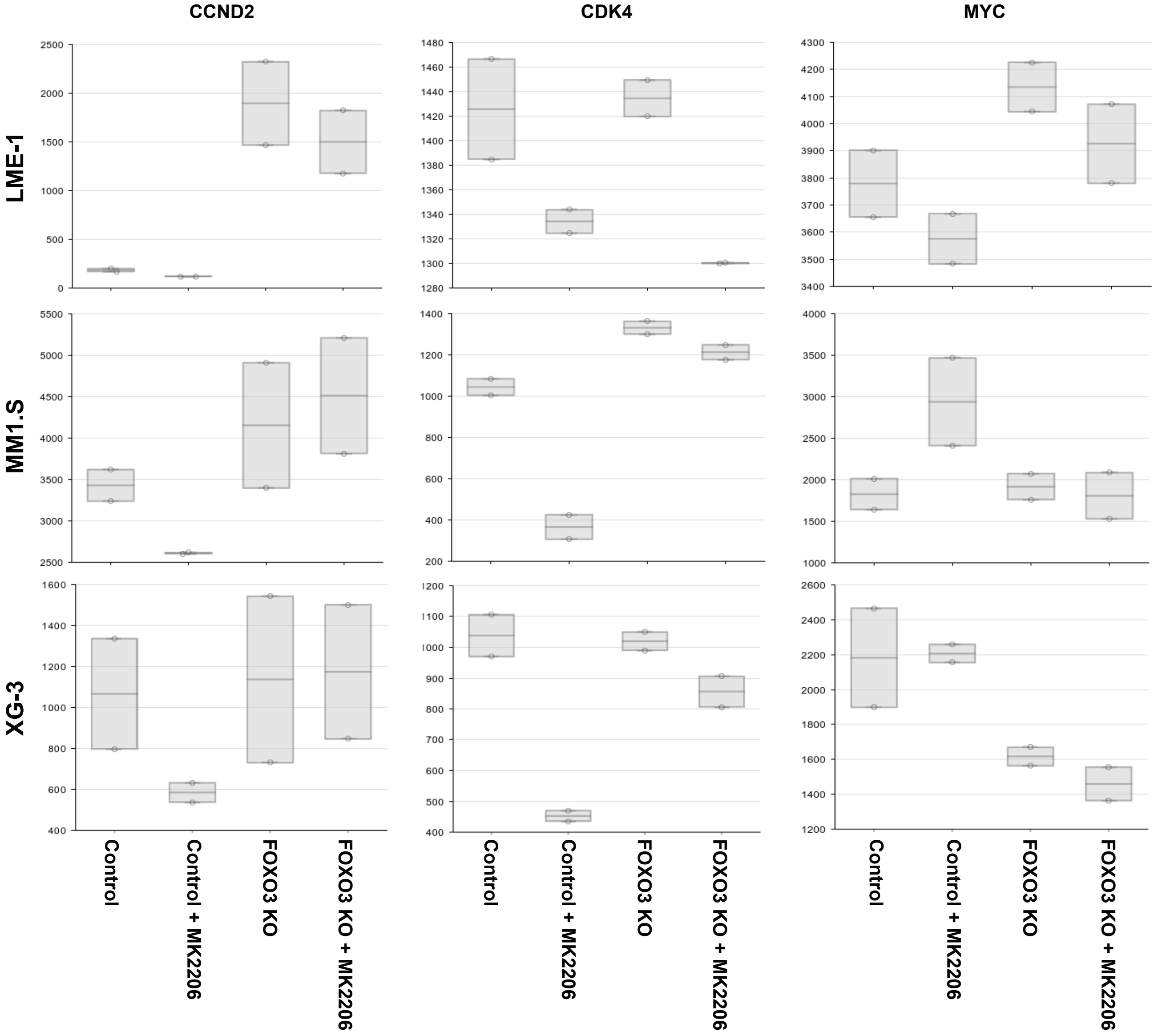


**SUPPLEMENTAL FIGURE 5. CCND2, CDK4 and MYC mRNA expression in HMCLs**
Independent LME-1 FOXO1 knockout clones (n=2), MM1.S FOXO3 knockout clones (n=2) and XG-3 FOXO3 knockout clones and their respective control clones (n=2) were treated overnight with 2.5 µM MK2206 AKT inhibitor, or left untreated, and subjected to gene expression profiling. Depicted are the normalized untransformed microarray gene expression levels as boxplots (each circle represents the mRNA levels of an independent clone) for CCND2, MYC and CDK4.

### SUPPLEMENTAL FIGURE 6

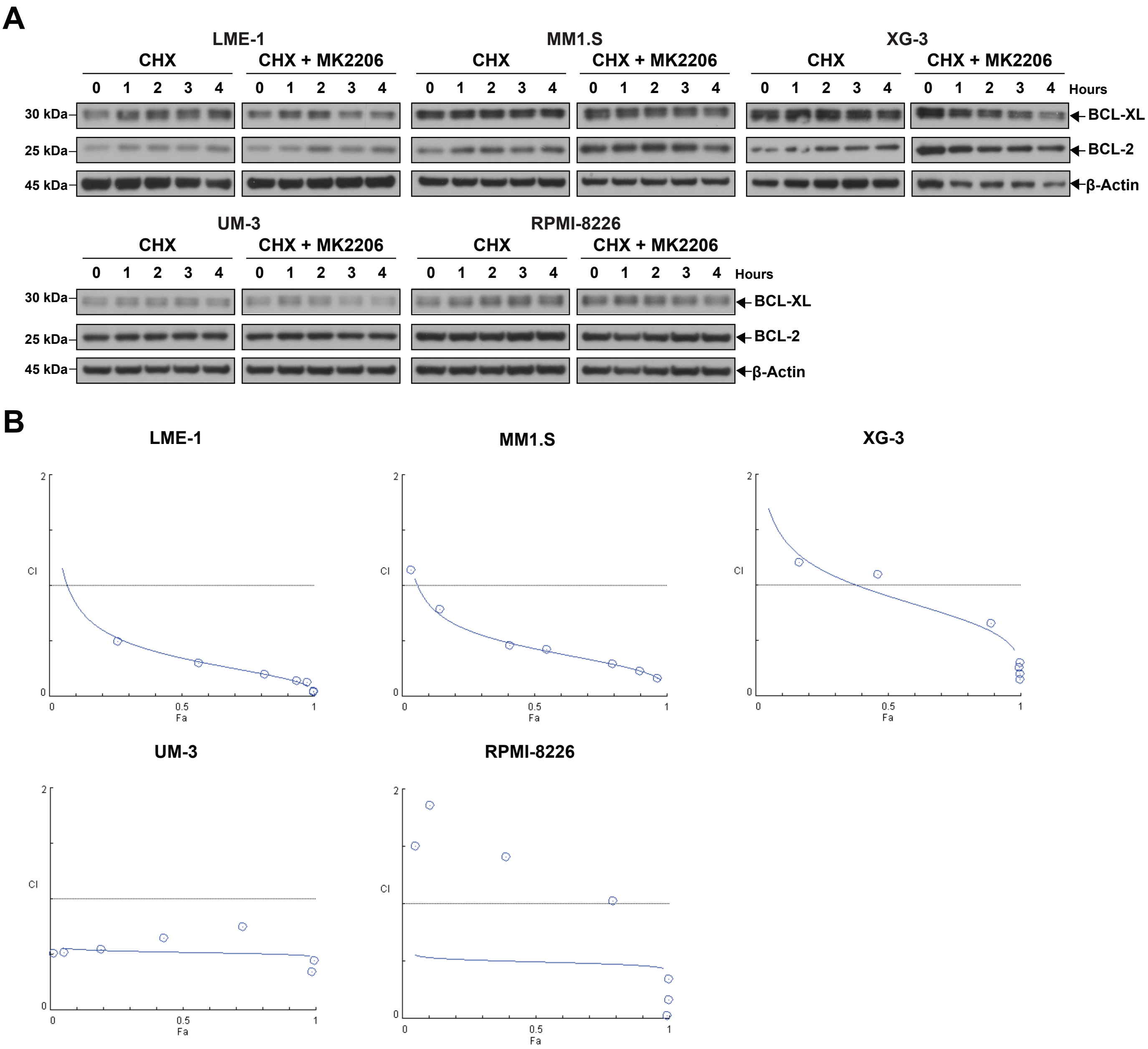


**SUPPLEMENTAL FIGURE 6. BCL2 and BCL-XL protein stability is not affected by AKT inhibition, Fa-CI plots of HMCLs treated with a combination of MK2206 and S63845**

**(A)** Immunoblot analysis of BCL2 and BCL-XL protein stability in HMCLs after cycloheximide (CHX) treatment, with or without a pretreatment of 2.5 μM MK2206 for 12 hours. Cells were treated with CHX as indicated by depicted time points. Immunoblots are from to the same experiment as Figure 6C, β-actin was used as a loading control. **(B)** Fa-CI plots showing the Chou-Talalay combination index (CI) as a function of the fraction of affected (Fa) after a combination treatment with MK2206 and S63845 at various concentrations. Horizontal lines represent CI = 1, signifying additive effects. Datapoints represent CI values at different drug concentrations. The Fa-CI plots directly correspond to the experiment depicted in Figure 6F
